# Examining models of modern human origins through the analysis of 43 fully sequenced human Y chromosomes

**DOI:** 10.1101/2023.11.09.566475

**Authors:** Shi Huang

## Abstract

Molecular studies have yielded two primary models for understanding the uniparental DNA phylogenetic trees of modern humans: the Recent Out of Africa (ROA) and the Recent Out of East Asia (ROE) models. These models differ in their underlying assumptions, particularly in relation to early stem haplotypes, even though they share many haplotype relationships. Leveraging the wealth of new genetic variants within the male-specific region unveiled through the comprehensive sequencing of 43 diverse human Y chromosomes, we here investigated the presence of shared variants among different haplotypes to determine which model better aligns with the genetic data. We were able to corroborate the existence of stem haplotypes specific to the ROE model, but not those exclusive to the ROA model. We found that A0b and A1a shared the most variants with each other, aligning with the A00A1a clade of the ROE model. Also, stem haplotypes specific to the ROE model showed the expected relationships, with A00A1a closest to B, AB closest to E, E closest to B, A, and C, ABDE closest to C. Our findings also revealed extensive variant sharing independent of common ancestry, consistent with the maximum genetic diversity theory underlying the ROE model but challenging the neutral theory behind the ROA model. These new genetic data lend robust support to the ROE model as the more accurate representation of modern human origins.

## Introduction

The origins of modern humans remain a subject of ongoing and intensive research. Molecular analyses of contemporary human populations have generated two distinct models to explain the evolutionary history of modern humans: the “Recent Out of Africa” (ROA) model and the “Recent Out of East Asia” (ROE) model. These models propose contrasting geographical locations as the point of origin for modern humans, with ROA suggesting Africa and ROE suggesting East Asia. The ROA model gained prominence in the 1980s, primarily based on mitochondrial DNA (mtDNA) studies (1, 2). On the other hand, the ROE model, also initially suggested by mtDNA studies in the 1980s (1), had been relatively overlooked until its recent independent rediscovery (3–6). Moreover, it has been revealed that all non-African Y chromosome haplotypes can be traced back to their origin in the southern part of East Asia (7).

The foundation of rooting uniparental DNAs in Africa rests on two fundamental assumptions: the molecular clock and the neutral theory (1, 2, 8, 9). Within this neutral framework, the “infinite site assumption” holds particular significance. This assumption posits that mutations occur only once in the course of evolutionary history, thereby excluding the possibility of back mutations and convergent mutations. It serves as a critical factor in deducing the derived alleles that are pivotal in shaping the phylogenetic tree topology of the ROA model (8, 10). However, the widely recognized downfall of the molecular clock hypothesis (11)—initially proposed as an *ad hoc* explanation for the genetic equidistance phenomenon—has effectively undermined the neutral theory, which was inspired by the molecular clock and supposed to explain or predict it (12). Quantifying constraint in the human mitochondrial genome has shown that most variants in mtDNA, including those in the non-coding region, are subject to selection (13). Direct experimental tests of the predictions of the neutral theory have disproven it (14). Strong evidence suggests that mutation saturation and natural selection are more prevalent than initially believed (3, 15–20). In the Y chromosome phylogenetic tree of the ROA model, numerous incompatible alleles have emerged, showcasing recurrent mutations (21). Ancient DNA has revealed common occurrence of convergent mutations in the human Y chromosome (22).

Genetic diversity patterns have continued to elude full explanation under the neutral theory (23), prompting the emergence of a novel framework in molecular evolution known as the “maximum genetic diversity theory” (17, 24–27). Central to this new theory is the intuitive notion, initially suggested by Fisher nearly a century ago, that mutations of a given size are less likely to be favorable in complex than simple organisms (28, 29). The most recent supporting evidence for this concept lies in the observation that lower species tend to harbor a greater number of common variants, which, however, are often detrimental in humans (30). In accordance with this new theory, and bolstered by a wealth of strong evidence, genetic diversities or distances are primarily observed to be at optimal saturation levels, rather than continually scaling with time, as assumed by the molecular clock and neutral theory. Upon mutation saturation, convergent evolution is very common where independent lineages share variants due to independent mutations and common selection.

The Y chromosome phylogenetic tree within the ROE model is constructed based on this novel theoretical framework, setting it apart from the ROA model in three significant ways (Fig. 1). Firstly, in ROE, haplotypes are defined by haplotype-specific alleles, irrespective of whether they are derived or result from back mutations. Secondly, the rooting in the ROE model is grounded in a straightforward premise: the original haplotype should be the one most commonly shared among individuals, as mutations leading to alternative haplotypes should be relatively rare occurrences (1). Finally, the haplotype representing the ancestral lineage is expected to exhibit a closer genetic relationship to ancient DNA samples from the time when modern humans initially appeared, in contrast to other contemporary haplotypes (3–5).

**Fig. 1.**
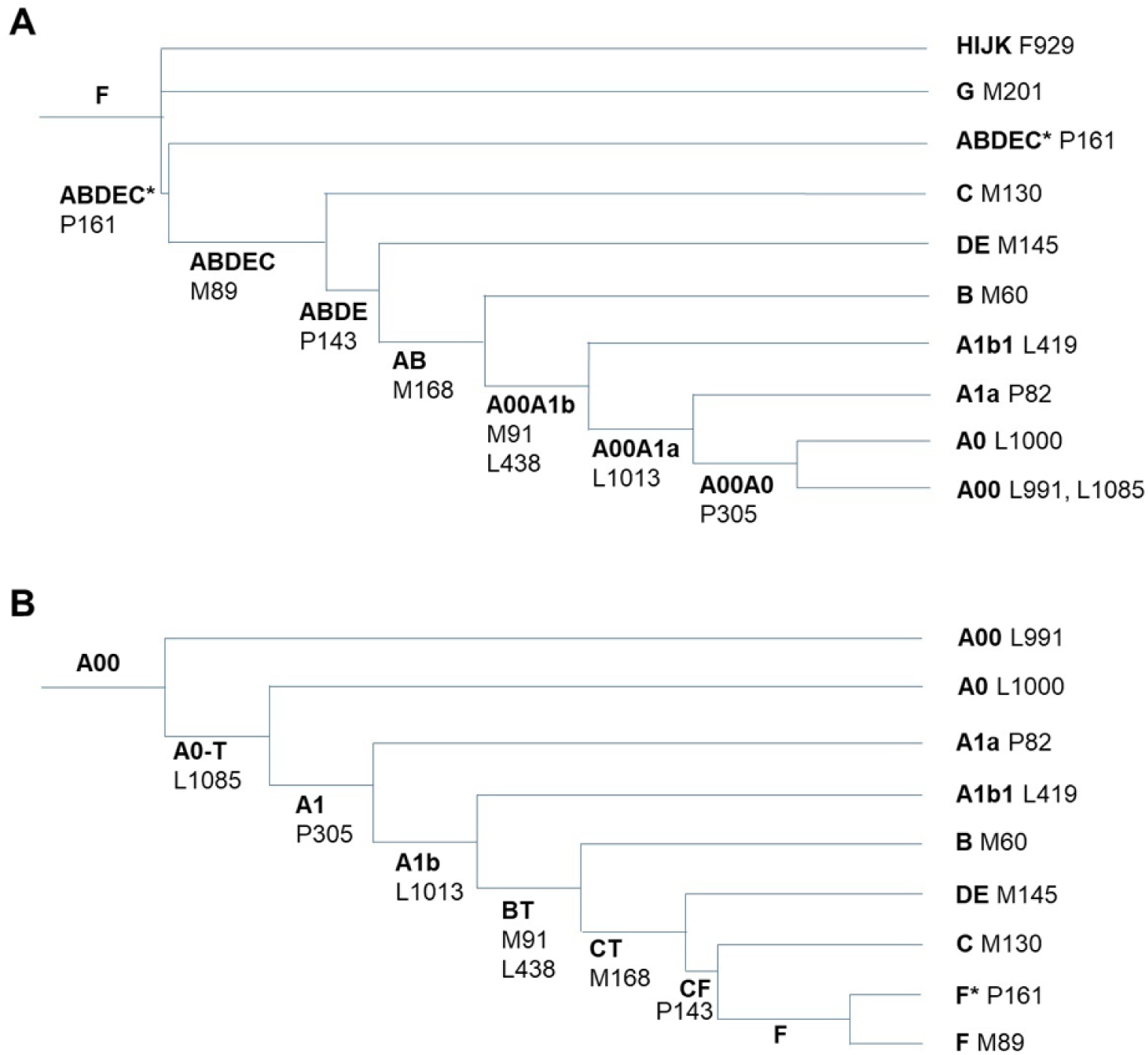
Y chromosome phylogenetic trees of modern humans. A. The Out of East Asia model. B. The Out of Africa model. Only major branches and representative SNVs are shown with branch lengths not to scale. The haplotype topologies for F subtypes under the ROA model, including G, H, I, J, and K, are consistent and identical between the ROA model and the ROE model. These haplotypes share the same genetic relationships and branching patterns in both models.

Both models, ROA and ROE, rely on assumptions that carry inherent uncertainties, necessitating independent testing to establish their validity. Human Y chromosomes have until recently only been partially sequenced (53.8%) and the phylogenetic tree of Y chromosome is built by using single nucleotide variants (SNVs, which are equivalent to SNP, with SNP often referring to common variants with frequency >0.01 whereas rare variants are sometimes referred as SNVs) located in the non-recombining Y region (NRY) or the male specific Y region (MSY). These variants are located within a 10 MB region amendable to short read sequencing (21). The human Y chromosome is 57 MB in length with 95% being MSY and 5% being the pseudoautosomal regions (PARs). The MSY region, unlike the PAR region, does not undergo recombination and is therefore informative to genealogy. A groundbreaking development is the recent achievement of complete sequencing for 43 human Y chromosomes, representing a wide array of major human groups or haplotypes (31). This comprehensive sequencing has unveiled a plethora of new SNVs and insertions or deletions (indels) located in the MSY region, which now offer a valuable resource for an independent examination of the human origin models of the paternal lineages.

In our study, we scrutinized these variants to ascertain which model finds stronger support in the context of this new variant data. Our results indicate that numerous variants are shared across different haplotypes, providing an informative pattern that sheds light on the relationships among these haplotypes. Importantly, this analysis validates the Y chromosome phylogenetic tree of the ROE model, offering compelling evidence in its favor.

## Methods and Materials

### Datasets

The data utilized in this study were sourced from Supplementary Tables S22, S24, and S25, as published in the paper by Hallast et al. in 2023 (31). These tables provide comprehensive listings of large and small size indels (Table S22 and S24) and SNVs (Table S25), along with the occurrences of these variants across the 43 Y chromosomes/samples. In our analysis, we combined the information from Table S22 and S24, which includes a total of 24,325 indels and 53,744 SNVs for our study. From these, we picked out variants from the MSY region for our studies based on their locations and Y-class designations as found in the source tables, including 16,679 indels and 34,764 SNVs.

### Calculation of the shared variant fraction between two haplotypes

For each haplotype/sample, we first compiled a list of SNVs or indels that are present within that specific haplotype/sample. We then employed custom scripts designed for this purpose to tally the count of shared genetic variants between the given haplotype/sample and all the other haplotypes/samples among the 43 Y chromosomes analyzed in our study. To quantify the extent of shared variants between two haplotypes, we computed the fraction of shared variants. For example, when the haplotype A0b (sample HG01890) was analyzed for the degree of sharing with each of the other 43 samples, the shared fraction was calculated by dividing the number of shared variants by the total number of variants in the A0b sample (6,793 SNVs or 3521 indels). These fraction values are listed in the A0b column of Table 2 and in Supplementary Tables S1 and S2. This approach allowed us to systematically assess the genetic relationships between the various haplotypes within our dataset.

### Calculation of the shared variant fraction between a stem haplotype and other haplotypes

A stem haplotype is ancestral and not directly observed in the current populations but is inferred to have existed in the past and is represented as the common ancestor of several modern haplotypes. The shared variant fraction between a stem haplotype and other haplotypes was calculated as follows, using the O stem haplotype as an example to illustrate our method. 1) Identification of SNVs or indels defining the O stem haplotype: SNVs or indels defining a stem haplotype, such as O, should be present in all terminal branch haplotypes. In the case of the O stem haplotype, there were one O1b sample and four O2a samples in the 43 Y chromosomes dataset. All SNVs or indels that were present in all five O samples were retrieved and designated as the O stem set. 2) Counting shared SNVs or indels: For a specific haplotype/sample, the number of shared SNVs or indels between that haplotype and the O stem set was counted. 3) Calculation of the shared fraction: The shared fraction for the O stem set was obtained by dividing the number of shared variants by the total number of variants in the O stem set. 4) Handling multiple samples: For haplotypes with multiple samples or terminal haplotypes represented in the 43 Y chromosomes dataset, such as haplotype E, the shared fraction between each individual E haplotype/sample and the O stem haplotype was independently calculated. These shared fractions for all E haplotypes/samples were then aggregated to calculate an average value. This average value was used to represent the shared fraction between the E haplotype as a whole and the O stem haplotype. This method allowed us to systematically assess and compare the shared genetic variants between the O stem haplotype and other haplotypes.

### Statistical methods

Standard methods including student’s t test and chi squared tests were used.

## Results

We used variants from the MSY region for our studies, including 16,679 indels and 34,764 SNVs located mostly in the ∼22 MB non-PAR euchromatin region (31). Within this dataset, we identified numerous variants that serve to uniquely characterize the non-controversial haplotypes shared by both competing models. As an example, the NO stem haplotype is defined by 75 SNVs and 9 indels, while the O stem haplotype is delineated by 106 SNVs and 15 indels (Table 1). Similarly, the RQ stem haplotype is defined by 6 SNVs. Furthermore, we uncovered a range of variants that define the stem haplotypes exclusive to the ROE model, including A00A1a, AB, ABDE, and ABDEC (Table 1 and Fig. 1A). An illustration of this is that A0b and A1a share 718 SNVs and 143 indels that are absent in other haplotypes/samples, supporting the A00A1a clade in the ROE model. In contrast, no variants were found that would support or uniquely define the ROA specific stem haplotype BT or CT (Table 1). The stem haplotype here refers to a haplotype that is found on an internal branch or node of a phylogenetic tree, representing a common ancestor from which several descendant haplotypes branch off. The terminal haplotype refers to a haplotype that is found at the tips (or leaves) of the phylogenetic tree. The results therefore support the ROE model and raise questions regarding the validity of the ROA model.

**Table 1.**
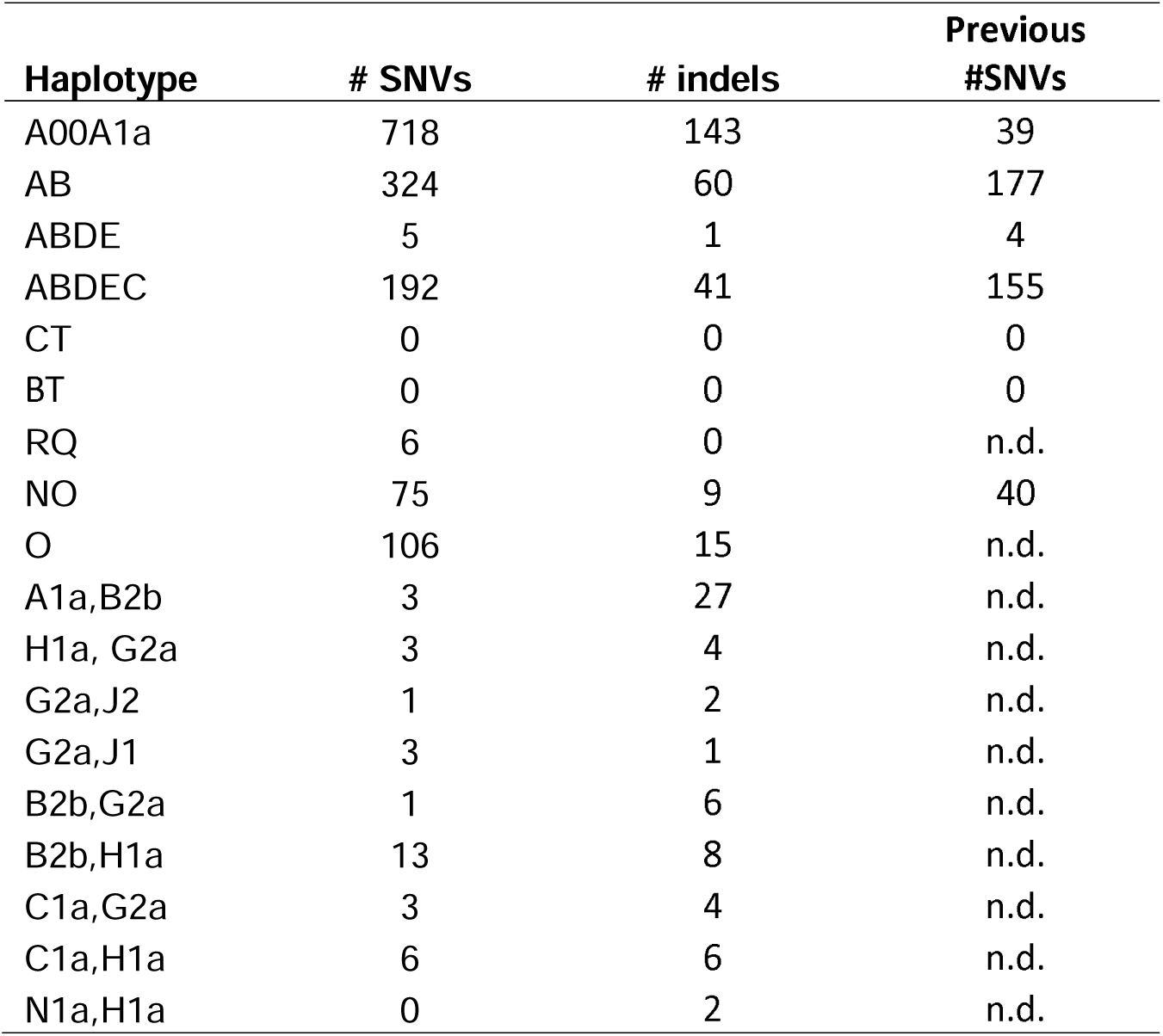
Number of variants uniquely found for each haplotype or combination of haplotypes. The listed haplotypes represent only a fraction of the complete set.

Notably, relatively to the number of variants found previously that define the ROE stem haplotypes, the full sequencing of 43 Y chromosomes has uncovered many new variants (Table 1). There are now 718 SNVs that define A00A1a, compared to previous 39 SNVs, 324 SNVs that define AB, compared to previous 177 SNVs, for example (22). Additionally, there are now a large number of indel variants that define these stem haplotypes (Table 1) but few such indels have previously been found. The results thus show that the full sequencing work has indeed as expected produced substantially more variants that are informative to phylogenetic analyses.

We also observed that many variants are present in two or more haplotypes that do not share a most recent common ancestor (MRCA), indicating convergent mutations. For example, H1a and G2a share 3 SNVs and 4 indels that are absent in other haplotypes/samples (H1a,G2a in Table 1). If these mutations had arisen in the MRCA of G and H, they would have been present in all H, I, J, and K haplotypes, as these 4 haplotypes share a more recent MRCA than with G. Other representative examples of sharing between two distantly related haplotypes include A1a,B2b; G2a,J2; G2a,J1; B2b,G2a; B2b,H1a; C1a,G2a; C1a,H1a;N1a,H1a (Table 1). These cases indicate the common occurrence of convergent mutations.

We systematically analyzed the pattern of shared variants among the 43 samples (Table 2, see Supplementary Table S1 for an expanded version of Table 2 containing all 43 samples and Supplementary Table S2 for the same analysis of indels). For example, among 6973 SNVs present in A0b, only 3275 are unique to A0b and the rest are shared in various degrees with each of the other haplotypes (8.3-39.6% shared) represented by the 43 samples (see A0b column in Table 2 and Supplementary Table S1). The results again indicate that convergent evolution may be common.

**Table 2.** Fractions of shared SNVs among selected haplotypes or samples. Each column represents the shared fractions between the sample listed at the first three rows and the samples listed in the first three columns. The bold numbers represent the shared SNV fraction value for a pair of haplotypes/samples, which exceeds the value obtained when each of the pair is compared to the 41 other haplotypes/samples individually. The samples were from the 1000 genomes project (ancestry groups are as indicated) and are listed based on Y chr phylogenetic relationships.

| Group | ID | Haplotype | # Total | # Unique | # Shared | ACB | GWD | LWK | YRI | GWD | GWD | MSL | ACB | LWK | LWK |
| --- | --- | --- | --- | --- | --- | --- | --- | --- | --- | --- | --- | --- | --- | --- | --- |
|  |  |  |  |  |  | HG01890 | HG02666 | NA19384 | NA19239 | HG03248 | HG02572 | HG03471 | HG02486 | NA19317 | NA19347 |
|  |  |  |  |  |  | A0b | A1a | B2b | E1a2a1a1a | E1a2b1a1a1 | E1a2b1a2 | E1b1a1a1a | E1b1a1a1a | E1b1a1a1a1c | E1b1a1a1a1c |
| ACB | HG01890 | A0b | 6973 | 3275 | 3698 |  | **0.465** | 0.470 | 0.454 | 0.522 | 0.468 | 0.490 | 0.484 | 0.484 | 0.511 |
| GWD | HG02666 | A1a | 5677 | 2486 | 3191 | **0.390** |  | 0.481 | 0.415 | 0.534 | 0.383 | 0.441 | 0.432 | 0.428 | 0.419 |
| LWK | NA19384 | B2b | 4126 | 1649 | 2477 | 0.278 | 0.391 |  | 0.421 | 0.550 | 0.404 | 0.462 | 0.459 | 0.449 | 0.447 |
| YRI | NA19239 | E1a2a1a1a | 3645 | 256 | 3389 | 0.237 | 0.295 | 0.372 |  | 0.892 | 0.739 | 0.594 | 0.558 | 0.542 | 0.522 |
| GWD | HG03248 | E1a2b1a1a1 | 2753 | 151 | 2602 | 0.206 | 0.317 | 0.367 | 0.674 |  | 0.600 | 0.502 | 0.489 | 0.478 | 0.459 |
| GWD | HG02572 | E1a2b1a2 | 4076 | 255 | 3821 | 0.274 | 0.301 | 0.399 | 0.827 | 0.888 |  | 0.611 | 0.588 | 0.580 | 0.577 |
| MSL | HG03471 | E1b1a1a1a1c | 3587 | 255 | 3332 | 0.252 | 0.268 | 0.402 | 0.584 | 0.654 | 0.538 |  | 0.845 | 0.808 | 0.782 |
| ACB | HG02486 | E1b1a1a1a1c1a1a | 3654 | 101 | 3553 | 0.254 | 0.319 | 0.407 | 0.559 | 0.649 | 0.527 | 0.861 |  | 0.890 | 0.863 |
| LWK | NA19317 | E1b1a1a1a1c1a1a3a1 | 3714 | 92 | 3622 | 0.258 | 0.336 | 0.404 | 0.552 | 0.645 | 0.528 | 0.837 | 0.905 |  | **0.911** |
| LWK | NA19347 | E1b1a1a1a1c1a1a3a1 | 3889 | 12 | 3877 | 0.285 | 0.342 | 0.421 | 0.557 | 0.649 | 0.551 | 0.848 | 0.919 | **0.954** |  |
| ESN | HG03371 | E1b1a1a1a1c1a1a3c1b | 3819 | 187 | 3632 | 0.253 | 0.324 | 0.413 | 0.567 | 0.646 | 0.534 | 0.876 | 0.930 | 0.903 | 0.881 |
| ACB | HG02011 | E1b1a1a1a1c1b | 3607 | 126 | 3481 | 0.252 | 0.299 | 0.410 | 0.578 | 0.644 | 0.544 | 0.882 | 0.868 | 0.872 | 0.833 |
| GWD | HG02717 | E1b1a1a1a1c2c3a2a | 3967 | 371 | 3596 | 0.265 | 0.293 | 0.407 | 0.580 | 0.657 | 0.535 | 0.865 | 0.872 | 0.848 | 0.816 |
| ACB | HG02554 | E1b1a1a1a2a1 | 3734 | 157 | 3577 | 0.260 | 0.313 | 0.403 | 0.570 | 0.642 | 0.520 | 0.847 | 0.857 | 0.844 | 0.806 |
| ESN | HG02953 | E1b1a1a1a2a1 | 3800 | 160 | 3640 | 0.260 | 0.325 | 0.423 | 0.579 | 0.660 | 0.551 | 0.883 | 0.905 | 0.864 | 0.842 |
| ASW | NA19705 | E1b1a1a1a2a1a3b1a1 | 3744 | 171 | 3573 | 0.256 | 0.325 | 0.400 | 0.558 | 0.633 | 0.534 | 0.853 | 0.871 | 0.866 | 0.823 |
| MSL | HG03065 | E1b1a1b1 | 4095 | 348 | 3747 | 0.244 | 0.303 | 0.398 | 0.610 | 0.638 | 0.582 | 0.785 | 0.786 | 0.780 | 0.755 |
| PUR | HG01109 | E1b1b1a1b1a | 3579 | 443 | 3136 | 0.223 | 0.301 | 0.408 | 0.540 | 0.632 | 0.525 | 0.572 | 0.570 | 0.555 | 0.545 |
| PUR | HG01106 | E1b1b1b1a1 | 3837 | 522 | 3315 | 0.256 | 0.276 | 0.373 | 0.607 | 0.640 | 0.570 | 0.616 | 0.610 | 0.591 | 0.597 |
| CLM | HG01457 | E1b1b1b2a1a4 | 3816 | 496 | 3320 | 0.244 | 0.285 | 0.400 | 0.603 | 0.617 | 0.562 | 0.634 | 0.626 | 0.623 | 0.607 |
| LWK | NA19331 | E1b1b1b2b2a1 | 3959 | 523 | 3436 | 0.251 | 0.304 | 0.362 | 0.617 | 0.655 | 0.576 | 0.601 | 0.569 | 0.556 | 0.578 |
| JPT | NA18989 | C1a | 5466 | 2522 | 2944 | 0.257 | 0.322 | 0.397 | 0.525 | 0.522 | 0.493 | 0.515 | 0.511 | 0.498 | 0.482 |
| MSL | HG03579 | G2a | 3318 | 966 | 2352 | 0.211 | 0.260 | 0.339 | 0.459 | 0.442 | 0.448 | 0.451 | 0.426 | 0.425 | 0.411 |
| BEB | HG03009 | H1a | 3161 | 1039 | 2122 | 0.220 | 0.233 | 0.343 | 0.431 | 0.468 | 0.401 | 0.471 | 0.444 | 0.430 | 0.423 |
| CLM | HG01258 | J1 | 3986 | 791 | 3195 | 0.218 | 0.264 | 0.358 | 0.487 | 0.456 | 0.486 | 0.470 | 0.452 | 0.447 | 0.444 |
| PJL | HG02492 | J2 | 2950 | 674 | 2276 | 0.189 | 0.296 | 0.345 | 0.358 | 0.463 | 0.333 | 0.387 | 0.380 | 0.361 | 0.354 |
| FIN | HG00358 | N1a | 2447 | 698 | 1749 | 0.185 | 0.237 | 0.310 | 0.321 | 0.426 | 0.306 | 0.345 | 0.337 | 0.328 | 0.324 |
| CHB | NA18534 | O1b | 3279 | 748 | 2531 | 0.203 | 0.246 | 0.310 | 0.453 | 0.424 | 0.447 | 0.424 | 0.402 | 0.394 | 0.384 |
| CHS | HG00673 | O2a2a1a1a | 2771 | 446 | 2325 | 0.197 | 0.228 | 0.325 | 0.396 | 0.433 | 0.371 | 0.367 | 0.359 | 0.349 | 0.373 |
| CHS | HG00621 | O2a2b1a1a1 | 2761 | 280 | 2481 | 0.212 | 0.243 | 0.304 | 0.412 | 0.441 | 0.381 | 0.430 | 0.402 | 0.397 | 0.406 |
| CHS | HG00512 | O2a2b1a2a1a | 3878 | 561 | 3317 | 0.231 | 0.243 | 0.348 | 0.483 | 0.438 | 0.478 | 0.451 | 0.429 | 0.432 | 0.458 |
| KHV | HG01596 | O2a1b1a1a1a1c | 3077 | 553 | 2524 | 0.217 | 0.259 | 0.323 | 0.408 | 0.442 | 0.373 | 0.419 | 0.413 | 0.413 | 0.400 |
| PEL | HG01928 | Q1b1a1a | 2481 | 207 | 2274 | 0.170 | 0.198 | 0.276 | 0.359 | 0.355 | 0.366 | 0.372 | 0.357 | 0.358 | 0.346 |
| PEL | HG01952 | Q1b1a1a | 2566 | 283 | 2283 | 0.162 | 0.217 | 0.256 | 0.378 | 0.339 | 0.357 | 0.355 | 0.326 | 0.324 | 0.314 |
| PJL | HG03492 | R1a1a1b2a1a2c | 2160 | 556 | 1604 | 0.139 | 0.168 | 0.206 | 0.349 | 0.292 | 0.292 | 0.309 | 0.281 | 0.265 | 0.257 |
| PUR | HG00731 | R1b1a1b1a1a2a5 | 1105 | 66 | 1039 | 0.091 | 0.138 | 0.136 | 0.216 | 0.186 | 0.186 | 0.215 | 0.192 | 0.184 | 0.184 |
| PUR | HG01243 | R1b1a1b1a1a2a5 | 1028 | 97 | 931 | 0.084 | 0.164 | 0.119 | 0.192 | 0.179 | 0.166 | 0.190 | 0.175 | 0.163 | 0.155 |
| MXL | NA19650 | R1b1a1b1a1a2b1a1 | 1135 | 124 | 1011 | 0.083 | 0.140 | 0.121 | 0.205 | 0.179 | 0.185 | 0.192 | 0.177 | 0.164 | 0.157 |
| TSI | NA20509 | R1b1a1b1a1a2b1a1 | 1314 | 91 | 1223 | 0.088 | 0.140 | 0.127 | 0.260 | 0.181 | 0.223 | 0.198 | 0.183 | 0.170 | 0.162 |
| CLM | HG01358 | R1b1a1b1a1a2b1a1 | 1319 | 121 | 1198 | 0.113 | 0.151 | 0.124 | 0.183 | 0.164 | 0.198 | 0.194 | 0.179 | 0.169 | 0.204 |
| GBR | HG00096 | R1b1a1b1a1a2c1a1f1a2 | 1216 | 189 | 1027 | 0.089 | 0.180 | 0.129 | 0.195 | 0.185 | 0.177 | 0.201 | 0.177 | 0.177 | 0.169 |
| IBS | HG01505 | R1b1a1b1a1a2c1a1a4b2d1 | 1322 | 132 | 1190 | 0.086 | 0.141 | 0.129 | 0.253 | 0.185 | 0.211 | 0.200 | 0.185 | 0.172 | 0.165 |
| ITU | HG03732 | R2 | 2019 | 597 | 1422 | 0.143 | 0.237 | 0.249 | 0.271 | 0.312 | 0.272 | 0.307 | 0.315 | 0.309 | 0.309 |

Haplotypes that are closely related, sharing a MRCA, are expected to exhibit a higher degree of variant sharing among them. This shared genetic material may arise from common ancestry and convergent mutations. Convergent mutations, for instance, may result from shared adaptations and two closely related groups with different Y haplotypes are expected to share more adaptations. To assess the phylogenetic relationships of different Y haplotypes, one can examine the extent of shared variants among them. If, among the 43 haplotypes or samples, haplotype 1 shares the highest fraction of variants with haplotype 2, and vice versa, these two haplotypes are considered the closest to each other or are likely to belong to the same clade. For example, sample NA19317 shared 95.4% of its variants with sample NA19347, a higher percentage than that observed between NA19317 and any other sample listed in the NA19317 column in Table 2. Similarly, NA19347 shared 91.1% of its variants with NA19317, which is greater than the percentage shared with any other sample listed in the NA19347 column in Table 2. These results suggest that these two samples are more closely related to each other than to any of the other samples studied here. Indeed, the two samples belong to the same clade, E1b1a1a1a1c1a1a3a1.

We determined the closest pairs among the 43 samples expected from the ROE model or the ROA model based on the known haplotype identity of each sample (Table 3). There are 12 pairs according the ROE model and 11 according to the ROA model. The two models differ in the expected pairing only with regard to the A0b-A1a pair, which are most related to each other in the ROE model belong to the A00A1a clade (Fig. 1a). In contrast, according to the ROA model, A1a, together with A1b, belongs to the A1 clade and A1a should be more closely related to haplotypes within the A1b clade, such as B, C, E, or F, among others (see Fig. 1B).

**Table 3.** Expected and observed pair of closely related samples within the 43 samples dataset.

| Group | ID | Haplotype | Expected closest relative |  | Observed |  |
| --- | --- | --- | --- | --- | --- | --- |
|  |  |  | OOEA | OOA | SNVs | Indels |
| ACB | HG01890 | A0b |  |  |  |  |
| GWD | HG02666 | A1a | HG01890 |  | yes | yes |
| LWK | NA19384 | B2b |  |  |  |  |
| YRI | NA19239 | E1a2a1a1a |  |  |  |  |
| GWD | HG03248 | E1a2b1a1a1 | HG02572 | same | no | no |
| GWD | HG02572 | E1a2b1a2 |  |  |  |  |
| MSL | HG03471 | E1b1a1a1a1c |  |  |  |  |
| ACB | HG02486 | E1b1a1a1a1c1a1a |  |  |  |  |
| LWK | NA19317 | E1b1a1a1a1c1a1a3a1 | NA19347 | same | yes | yes |
| LWK | NA19347 | E1b1a1a1a1c1a1a3a1 |  |  |  |  |
| ESN | HG03371 | E1b1a1a1a1c1a1a3c1b |  |  |  |  |
| ACB | HG02011 | E1b1a1a1a1c1b |  |  |  |  |
| GWD | HG02717 | E1b1a1a1a1c2c3a2a | HG02011 or HG02554/3 | same | yes | no |
| ACB | HG02554 | E1b1a1a1a2a1 |  |  |  |  |
| ESN | HG02953 | E1b1a1a1a2a1 |  |  |  |  |
| ASW | NA19705 | E1b1a1a1a2a1a3b1a1 | HG02953 or HG02554 | same | yes | no |
| MSL | HG03065 | E1b1a1b1 |  |  |  |  |
| PUR | HG01109 | E1b1b1a1b1a |  |  |  |  |
| PUR | HG01106 | E1b1b1b1a1 |  |  |  |  |
| CLM | HG01457 | E1b1b1b2a1a4 | NA19331 | same | no | no |
| LWK | NA19331 | E1b1b1b2b2a1 |  |  |  |  |
| JPT | NA18989 | C1a |  |  |  |  |
| MSL | HG03579 | G2a |  |  |  |  |
| BEB | HG03009 | H1a |  |  |  |  |
| CLM | HG01258 | J1 | HG02492 | same | no | no |
| PJL | HG02492 | J2 |  |  |  |  |
| FIN | HG00358 | N1a |  |  |  |  |
| CHB | NA18534 | O1b |  |  |  |  |
| CHS | HG00673 | O2a2a1a1a |  |  |  |  |
| CHS | HG00621 | O2a2b1a1a1 |  |  |  |  |
| CHS | HG00512 | O2a2b1a2a1a | HG00621 or HG00596 | same | yes | yes |
| KHV | HG01596 | O2a1b1a1a1a1c |  |  |  |  |
| PEL | HG01928 | Q1b1a1a | HG01952 | same | yes | yes |
| PEL | HG01952 | Q1b1a1a |  |  |  |  |
| PJL | HG03492 | R1a1a1b2a1a2c |  |  |  |  |
| PUR | HG00731 | R1b1a1b1a1a2a5 | HG01243 | same | no | no |
| PUR | HG01243 | R1b1a1b1a1a2a5 |  |  |  |  |
| MXL | NA19650 | R1b1a1b1a1a2b1a1 |  |  |  |  |
| TSI | NA20509 | R1b1a1b1a1a2b1a1 | NA19650 or HG01358 | same | no | no |
| CLM | HG01358 | R1b1a1b1a1a2b1a1 |  |  |  |  |
| GBR | HG00096 | R1b1a1b1a1a2c1a1f1a2 | HG01505 | same | no | yes |
| IBS | HG01505 | R1b1a1b1a1a2c1a4b2d1 |  |  |  |  |
| ITU | HG03732 | R2 |  |  |  |  |

Our approach identified 6 of 12 pairs in the ROE model that emerged as each other’s closest relatives based on the shared SNV fractions among the 43 samples (Table 2 and Supplementary Table S1). Remarkably, the A0b and A1a pair was among the 6 identified.

One possibility might explain that not all expected pairs could be identified by our approach here. Two haplotypes may have diverged for very long time such as J1 and J2 and may each accumulate many independent mutations. Convergent evolution could result in each being more related to a haplotype from a different clade. This could also similarly account for recently diverged haplotypes not sharing the most with each other; they may have encountered strong adaptive evolution with a haplotype from a distantly related clade.

While not all closely related pairs of haplotypes could be identified as such by our approach, it is important to note that there were not any pair of unrelated or distantly related haplotypes that were mis-identified as the most closely related by our method. For instance, G2a/HG03579 shared the most SNVs (57.7%) with the distantly J1/HG01258, while J1/HG01258 shared the most (51.8%) with the distantly related E1b1a1b1E/HG03065, rather than with G2a/HG03579 (Supplementary Table S1). Given this observation, it is highly unlikely for the A0b-A1a pair, otherwise distantly related according to the ROA model, to represent an exception to the pattern that only closely related haplotypes could be identified as such by our approach.

We also performed the same analysis by using indel variants and obtained consistent results (Supplemental Table S2). Together, these findings strongly support the A00A1a clade within the ROE model, which encompasses A0b and A1a as terminal branches. The data therefore once again support the ROE model and challenge the ROA model.

According to the ROA model, the expected topological relationship between A1a and A0b is akin to that between B and A1b1 (or A1a) or between E and B (as depicted in Fig. 1B). Thus, if the ROA model predicts that A1a should exhibit the highest degree of variant sharing with A0b, it would logically follow that E should share the most variants with B. To put this prediction to the test, we conducted a thorough analysis of the sharing patterns among all individual haplotypes and samples, as presented in Fig. 2. We used the average value of the shared fractions of all individual E haplotypes or samples to represent the value of E. The results revealed a significant disparity from the expected outcomes under the ROA model. E was found to share more variants with C1a than with B (Fig. 2A). Similar analysis using indel variants revealed consistent results (Fig. 2B). Therefore, the close relationship observed between A0b and A1a contradicts the ROA model.

**Fig. 2.**
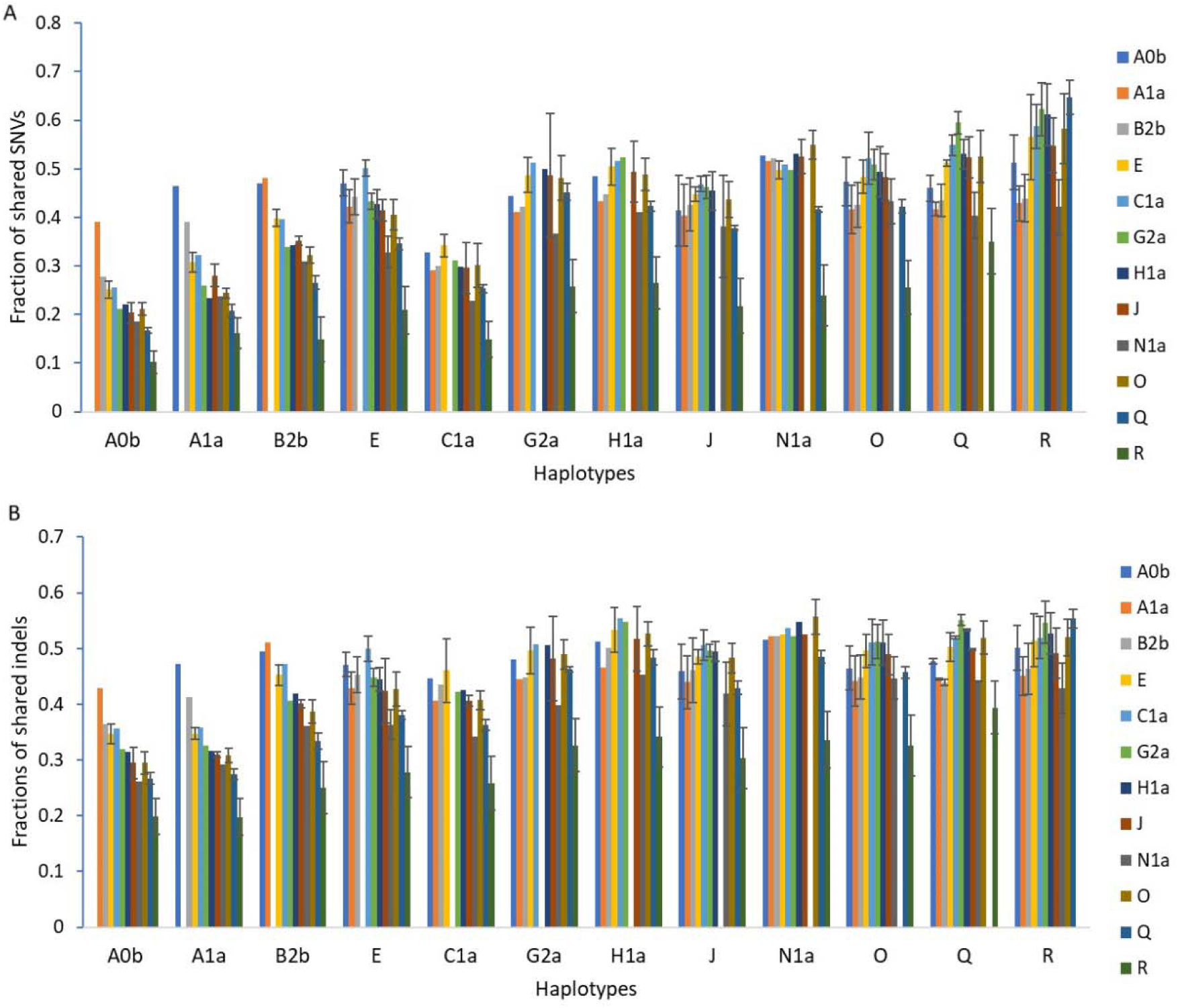
Fraction of shared variants among individual terminal haplotypes. A. Fraction of shared SNVs. B. Fraction of shared indels. In the case of haplotypes like E, J, O, Q, and R that have more than one individual terminal haplotype/sample represented in the dataset of 43 chromosomes, the average value of all samples under a haplotype is shown with error bars representing the standard deviation of the data. The information used to generate the figures were extracted from Supplementary Table S1 and S2.

The approach of inferring close relationships based on the level of shared variants offers a fresh perspective for evaluating the two competing models of human origins. We next focused on variants that define a stem haplotype. These variants are present in all individual terminal haplotypes belonging to the respective stem haplotype. Such stem-defining variants are more informative to the relationship between a stem haplotype and another haplotype compared to the variants found in a terminal haplotype. We examined whether the stem haplotypes exclusive to the ROE model exhibited the anticipated relationships. As depicted in Fig. 3, the stem haplotype A00A1a of the ROE model shared the most variants with B2b, a finding that aligns with the presence of the AB stem haplotype. The AB stem haplotype in the ROE model shared the most variants with E, while the stem haplotype E shared the most with A and B (except in the case of indels where E shared more with C than A00A1a, Fig. 3B), all consistent with the existence of the ABDE stem haplotype. Notably, the ABDE stem haplotype shared the most variants with C, reinforcing the presence of the ABDEC stem haplotype. Additionally, F was found to share variants nearly equally with A, B, E, and C, which is in line with expectations derived from the ROE model. However, F should be closer to C than to A, B, and E if the CF clade in the ROA model is real. These results lend substantial support to the actual existence of all the stem haplotypes unique to the ROE model. By contrast, the predictions derived from the ROA model do not align with the observed relationships in Fig. 3. Therefore, these findings further challenge the ROA model.

**Fig. 3.**
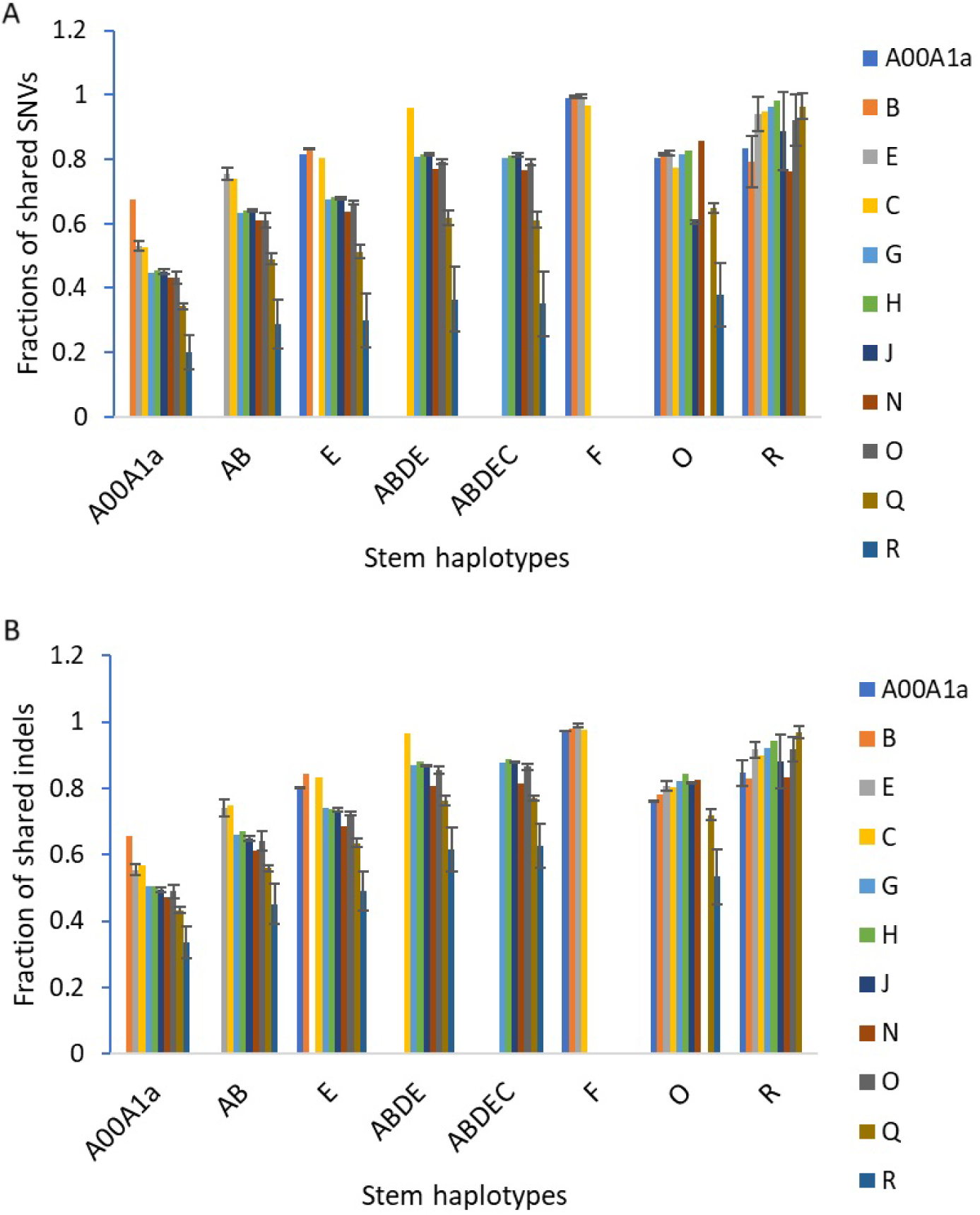
Fraction of shared variants between a stem haplotype in the ROE model and other individual terminal haplotypes in the 43 human Y chromosomes. A. Fraction of shared SNVs. B. Fraction of shared indels. For haplotypes such as A, E, J, O, Q, and R that have more than one individual terminal haplotype/sample represented in the 43 chromosomes, the average value of all samples under a haplotype is shown with error bars representing the standard deviation.

All the stem haplotypes within the ROE model that are common in African populations, including A00A1a, AB, ABDE, and ABDEC, are anticipated to exhibit equal genetic relatedness to all F subtypes such as G, H, J, N, O, Q, and R. The observations presented in Fig. 3 is largely in line with this expectation where G, H, and J shared similar number of variants with these African stem haplotypes. However, haplotype within the K2 clade (R, Q, O, and N) shared lower number of variants with these African stem haplotypes compared to G, H, or J. This can likely be attributed to the fact that the reference genome used for variant calling (GRCh38 Y) corresponds to the R1b haplotype, which would inherently result in a reduced number of variants called for R and its closely related Q, O, or N haplotype, as demonstrated in Table 2 and Supplementary Table S1-S2 where the total number of variants called for each haplotype or sample are listed. Some of the variants that were not called could include some shared variants with the African stem haplotypes.

We extended our analysis to assess the stem haplotypes exclusive to the ROA model, including BT, CT, and CF. According to the ROA model’s predictions, BT should exhibit the closest relationship to A1a, CT should be most closely related to B, and CF should have the strongest affinity with E. However, our findings deviate from these expectations. Our results indicated that BT is similarly related to A0b and A1a, or shares slightly fewer variants with E1a than with A0b. CT is similarly related to B2b, A1a, and A0b, while CF is similarly related to E, B2b, A1a, and A0b (Fig. 4). These results raise questions about the validity of the stem haplotypes unique to the ROA model, suggesting that they may not correspond to actual genetic relationships.

**Fig. 4.**
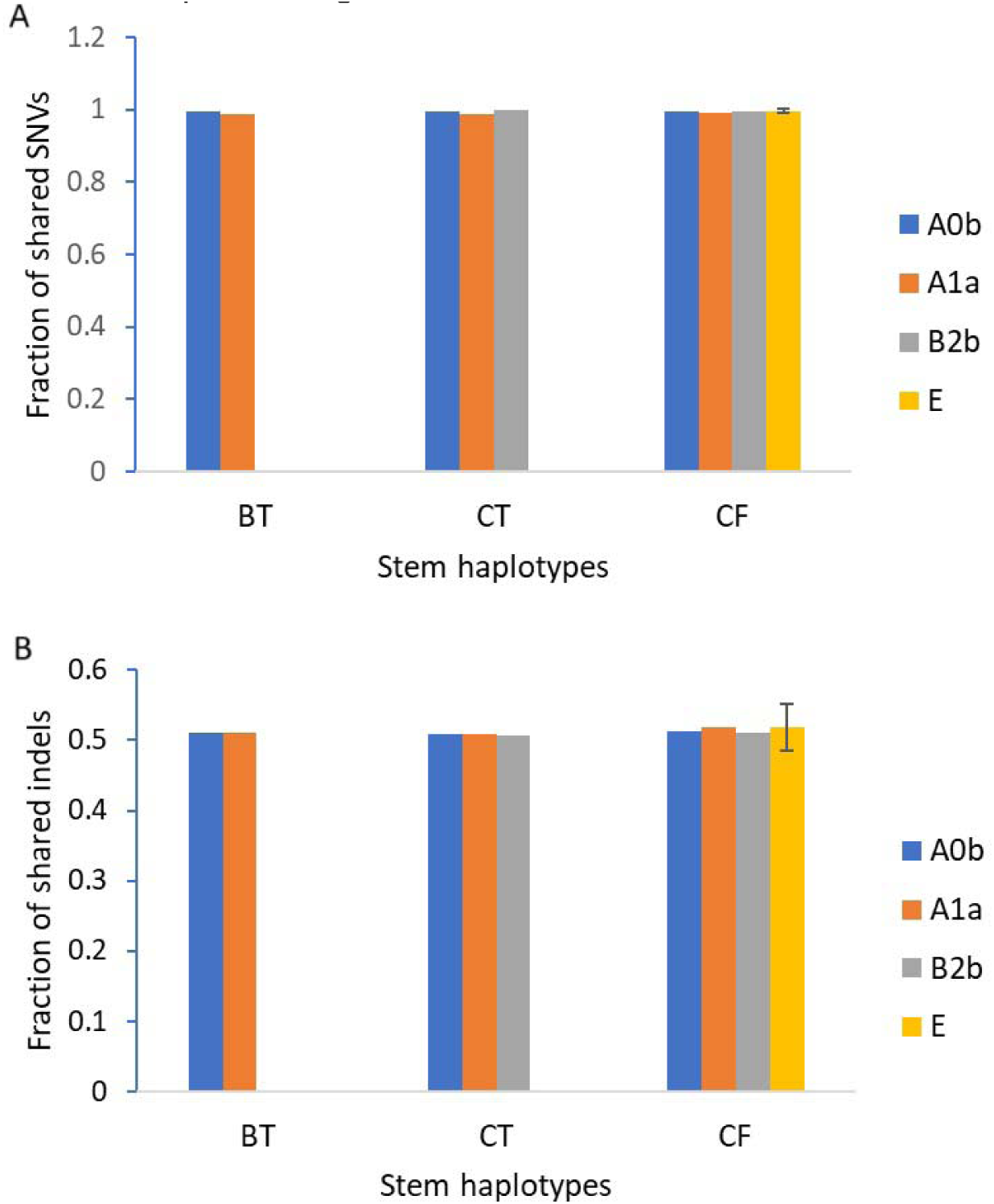
Fraction of shared variants between a stem haplotype in the ROA model and other individual terminal haplotypes in the 43 human Y chromosomes. A. Fraction of shared SNVs. B. Fraction of shared indels. For haplotypes such as E that have more than one individual terminal haplotype/sample represented in the 43 chromosomes, the average value of all samples under a stem haplotype is shown with error bars representing the standard deviation.

Finally, we studied the proportion of variant sharing between two haplotypes that is due to convergent evolution. As shown in Fig. 4, for the O1b (NA18534) and O2a (HG00673) pair, O1b (NA18534) has 3279 SNVs and shared 1651 with O2a (HG00673) and 1254 with all four O2a samples in the dataset. So, the number of shared variants between the O1b and O2a samples that are not due to common ancestry is 397 (1651–1254). The proportion of shared variants due to convergent evolution is thus 0.24 (397/3279). Similarly, the proportion of convergent evolution for the E1a (NA19239) and E1b (HG03471) pair is calculated to be 0.31, and for the R1a (HG03492) and R1b (HG00731) pair is calculated to be 0.48 (Fig. 5). Analyses using indel variants showed similar results but the proportion of convergent evolution is higher compared to SNVs (Fig. 5). These data indicate that a substantial fraction of shared variants between any two haplotypes is due to convergent evolution.

**Fig. 5.**
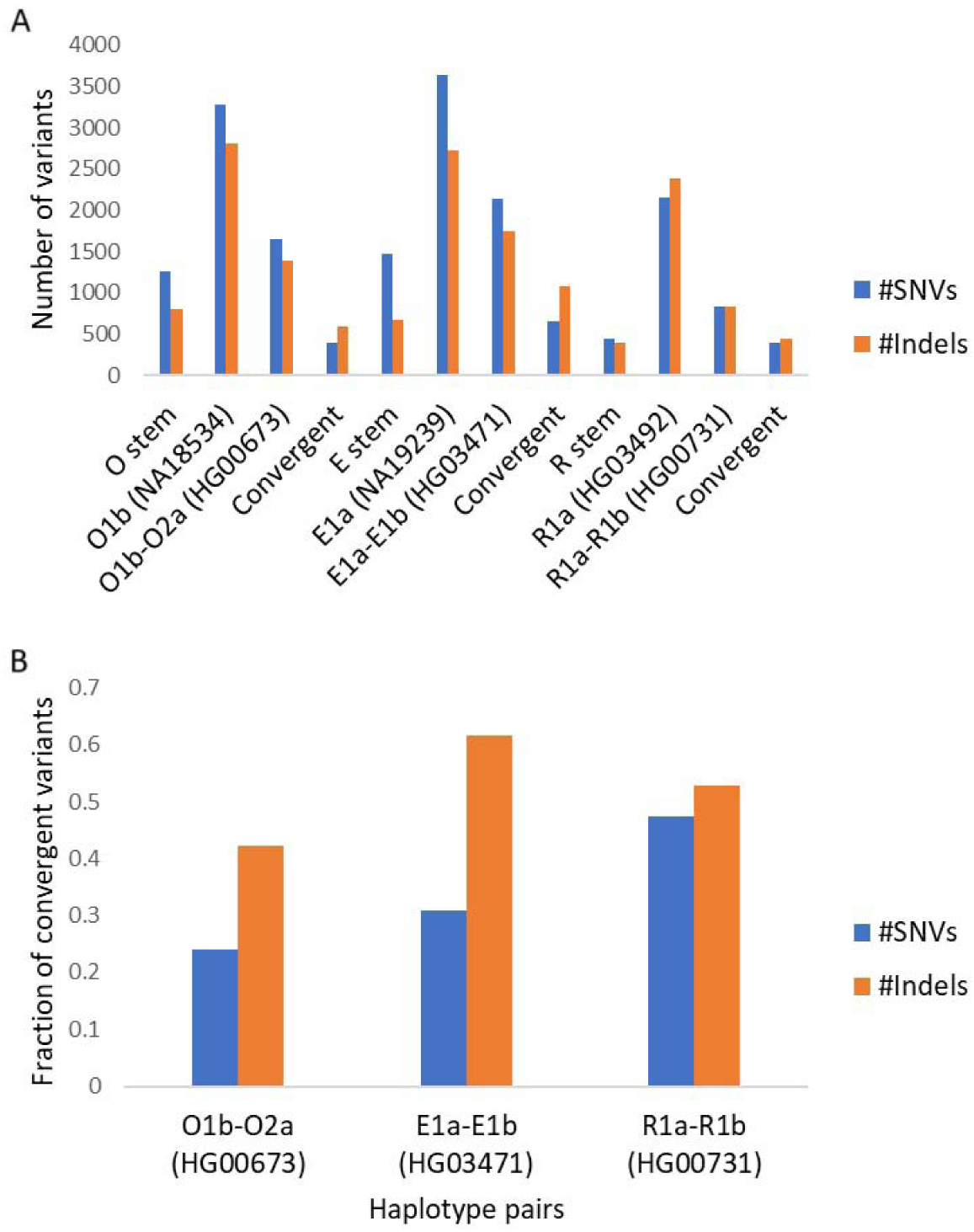
Number and fraction of shared variants due to convergent evolution in selected haplotypes. A. Number of variants. B. Fraction of shared variants due to convergent evolution.

## Discussions

Within the ROE model, a well-defined hierarchy is observed, where A0b and A1a fall within the A00A1a clade, A00A1a resides within the AB clade, AB is within ABDE clade, E is encompassed within the ABDE, ABDE is nested within the ABDEC clade, and C finds its place within the ABDEC clade. Remarkably, the anticipated variant sharing patterns are in good alignment with the actual observations we have presented in our study. These findings offer compelling support for the representation of the genetic relationships among these haplotypes as offered by the ROE model.

Our study presents three observations that pose significant challenges to the ROA model. First, in contrast to the predictions by the ROA model, we found that A1a exhibited the closest relationship with A0b. However, this observation was not paralleled by the expected proximity of E to B as prescribed by the ROA model, if one were to consider the possibility of this observation being compatible with the ROA model. Second, according to the ROA model, the E stem haplotype should be more closely related to C than to B, and the F stem haplotype should have a closer genetic affinity with C than with E. Our study did not align with these expectations. Lastly, the stem haplotypes unique to the ROA model, namely BT, CT, and CF, all failed to exhibit the anticipated sharing patterns. These observations collectively cast doubt on the explanatory power of the ROA model to account for the genetic relationships we observed in our study.

Influenced by the neutral theory, sharing of variants between any two haplotypes of Y chromosome is previously thought to be all due to common ancestry. Our findings here however showed that a substantial fraction of shared variants may be due to convergent evolution (Fig. 5). Previous studies only used variants located in certain Y chromosomal regions that can be easily identified by short read sequencing and did not observe significant convergent evolution (21).

We have also employed ancient DNA analysis to provide independent validation of the ROE model, as detailed in our previous work (22). The ancient DNA study also revealed the common occurrence of convergent evolution. Ancient samples only show mutations in a portion of the loci defining a haplotype, whether it is a basal haplotype or a terminal haplotype. This implies that the diversification of terminal haplotypes had actually occurred before the basal haplotype had completely mutated all relevant loci, contrary to what the neutral theory predicts. This also suggests that the process of the basal haplotype transitioning from partial to complete mutation of all loci occurs simultaneously with the full formation of the terminal haplotype, necessitating the involvement of convergent mutations. This is akin to the growth of a living tree, where the development of leaves or branches occurs in sync with the growth of the trunk.

The ROE model is also strongly supported by the recent discoveries in human fossils. The Yunxian crania from Hubei China, dated to 1 million years ago, are more modern than *H. erectus* (reduced alveolar prognathism and no sagittal crest) and are likely on the lineage leading to modern humans (32–36). The Hualongdong skull HLD6 from Anhui China at 300,000 years ago also has features of modern humans such as a weak chin, high eye socket, flat face, a pronounced mental trigone, and a weak alveolar prognathism (37, 38). The Dingcun teeth from Shanxi China are ∼270,000 years old and show modern features that are not found in Southern Africa fossils older than 100,000 years (39). It should be noted that different skull traits are not equally informative when identifying modern human forms. Alveolar prognathism appears to be a highly informative trait, yet it has long been somewhat overlooked. Most human fossils from Africa, regardless of age, including Jebel Irhoud at 300,000 year ago (40), as well as a high fraction of present-day sub-Sahara Africans, have alveolar prognathism. If humans with some modern features had been living in East Asia at 1 million or 300,000 years ago, it would be expected—barring any unforeseen random events—that modern humans with the most complete modern traits would also have lived in East Asia during the period when fully modern humans first emerged, i.e., between 100,000 and 50,000 years ago. Thus, the ROE model is well aligned with the fossil record.

The ROE model is more consistent with first principle logics. The group with the leading cognitive abilities and other phenotypic traits should be the first to cross the threshold into modern human status and maintain that status, barring any unexpected events. Compared to other ancient human groups, the first group to become modern humans should have a genome most distant from chimpanzees and maintain this distinction to the present. The first population to become modern humans should have lower maximum genetic diversity and maintain this low diversity unless disrupted by unexpected events.

The regions suitable for the evolution of *Homo erectus* into modern humans should also be suitable for the continued evolution of modern humans. Conversely, regions where current populations are less advanced should be less likely to have been the cradle of early modern humans. The ROA hypothesis contains an inherent contradiction: it assumes that, while Africa was more conducive to human progress between 2 million and 50,000 years ago, Eurasia became more suitable only in the last 50,000 years. As a result, modern humans in Eurasia are genetically further removed from the great apes. (41).

The complex and diverse environment of East Asia provides moderate challenges suitable for progressive human evolution (42). Tropical regions have more pathogens, which require populations living there to have stronger immunity. This increased immunity, however, may not be conducive to reducing genetic diversity. As Gage noted about the higher genetic diversity of chimpanzees compared to humans, “Humans might rely less on biological variation than cultural and cognitive evolution—in the form of medicine and engineering, for example—to adapt to changing environments.” (43). By this sound reasoning, human populations with relatively lower genetic diversity may depend more on cultural and cognitive evolution than on biological evolution. Existing research on culture and cognitive abilities appears to support this intuitive perspective.

A model should be self-consistent. Yet, the ROA model has a major self-contradiction. For the first seven major haplotype splits in the Y chromosome tree (Figure 1B, from A00 to F), the population associated with each newly derived haplotype is consistently much larger than the ancestral population. For example, the A0-T population is orders of magnitude greater than the ancestral A00 population, the BT population is orders of magnitude greater than the ancestral A population, and so on, with the F population being much larger than the C population at the seventh split, implying a fitness advantage for each of the newly formed haplotypes, therefore contradicting the neutral assumption required for building the tree in the first place. The same issue also exists for the mtDNA tree of the ROA model.

If all haplotypes are equally fit, as assumed by the neutral theory, the original ancestral haplotype of Y chromosome or mtDNA should be the most abundant and widespread. This is because mutations leading to other haplotypes are rare events, occurring only in a fraction of the entire population. Yet, the ancestral haplotype in the ROA model, A00 for Y chromosome and L0 for mtDNA, is the rarest. In contrast, the ancestral type in the ROE model, F for Y chromosome and R for mtDNA, is the most abundant (Figure 1A).

The ROE theory does not necessitate concealing any truths, whereas the ROA theory often overlooks certain scientific realities, such as the differences in genetic and phenotypic distances between various human populations and great apes (41). Furthermore, the ROA advocates tend to downplay the foundational role of the neutral theory in the ROA model and the considerable uncertainty or controversy surrounding that theory.

The ROA theory and its associated neutral theory provide only a limited understanding of human uniqueness in terms of DNA, lacking insights into the differences between humans and lower animals, as well as the distinctions between modern and archaic humans. If one lacks a correct understanding of the specific genomic features of humans, it would be difficult to infer human evolutionary processes based on DNA data.

The results presented in our study offer yet another strong support for the ROE model. Future molecular tests to further assess these Y chromosome models may concentrate on obtaining more comprehensive sequencing data from additional ancient and contemporary human Y chromosomes, with particular emphasis on populations from Africa and East Asia. Expanding the scope of investigations to encompass a wider array of genetic data will contribute to a more comprehensive and refined understanding of human origins.

## Acknowledgements

This research was conducted without the direct support of any specific research grants.

